# Coiled coils 9-to-5: Rational *de novo* design of α-helical barrels with tunable oligomeric states

**DOI:** 10.1101/2021.01.20.427391

**Authors:** William M. Dawson, Freddie J.O. Martin, Guto G. Rhys, Kathryn L. Shelley, R. Leo Brady, Derek N. Woolfson

## Abstract

The rational design of linear peptides that assemble controllably and predictably in water is challenging. Sequences must encode unique target structures and avoid alternative states. However, the stabilizing and discriminating non-covalent forces available are weak in water. Nonetheless, for α-helical coiled-coil assemblies considerable progress has been made in rational *de novo* design. In these, sequence repeats of nominally hydrophobic (***h***) and polar (***p***) residues, ***hpphppp***, direct the assembly of amphipathic helices into dimeric to tetrameric bundles. Expanding this pattern to ***hpphhph*** can produce larger α-helical barrels. Here, we show that pentamers to nonamers are achieved simply by varying the residue at one of these ***h*** sites. In L/I-K-E-I-A-x-Z repeats, decreasing the size of Z from threonine to serine to alanine to glycine gives progressively larger oligomers. X-ray crystal structures of the resulting α-helical barrels rationalize this: side chains at Z point directly into the helical interfaces, and smaller residues allow closer helix contacts and larger assemblies.

---

Most commonly, natural coiled-coil (CC) peptides form dimers, trimers and tetramers with consolidated hydrophobic cores.^1–2^ Control over oligomeric state is achieved by different combinations of mainly isoleucine (Ile, I) and leucine (Leu, L) residues in the core.^3–4^ Larger oligomers are rare in nature.^5–6^ Interestingly, some of these larger structures are α-helical barrels (αHBs) with accessible central channels making them appealing scaffolds for functional design, *e.g.* binding, catalysis, delivery and transport.^7–13^ Variants of a natural dimer and *de novo* tetramer serendipitously form heptameric and hexameric αHBs, respectively.^14–15^ To automate the design of αHBs, we have developed computational-design tools to deliver 5-, 6- or 7-helix αHBs.^16^ These oligomers can be rationalized retrospectively to advance further sequence-to-structure relationships for CC design.

Most αHBs are Type-2 CCs based on ***hpphhph*** sequence repeats, labelled ***abcdefg*** (Figure. 1).^17^ Typically, αHBs have L/IxxIAxA repeats; *i.e., **a*** = Leu or Ile and ***d*** = Ile. β-Branched residues at ***d*** are particularly important for maintaining open αHBs.^18^ The hexameric and heptameric αHBs (CC-Hex2 and CC-Hept, systematically named CC-Type2-(S_g_L_a_I_d_)_4_ and CC-Type2-(A_g_L_a_I_d_)_4_) have ***a*** = Leu, ***d*** = Ile and ***e*** = alanine (Ala, A), but differ at ***g***, which is Ala in the heptamer and the slightly larger serine (Ser, S) in the hexamer (Table 1). Another variant, CC-Pent (CC-Type2-(I_g_L_a_I_d_E_e_)_4_) has Ile at ***g***, although it differs from the other examples having ***e*** = glutamic acid (Glu, E).^16^ A second series with all-Ile cores (***a*** = ***d*** = Ile) has been characterized.^16,18^ In these, another hexamer, CC-Hex3 (CC-Type2-(S_g_I_a_I_d_)_4_), follows the design rules above (Table 1);^16^ and a peptide with Ala at ***g*** (CC-Type2-(A_g_I_a_I_d_)_4_) forms an octamer when crystallized in the presence of isopropanol.^18^

**Figure 1.**
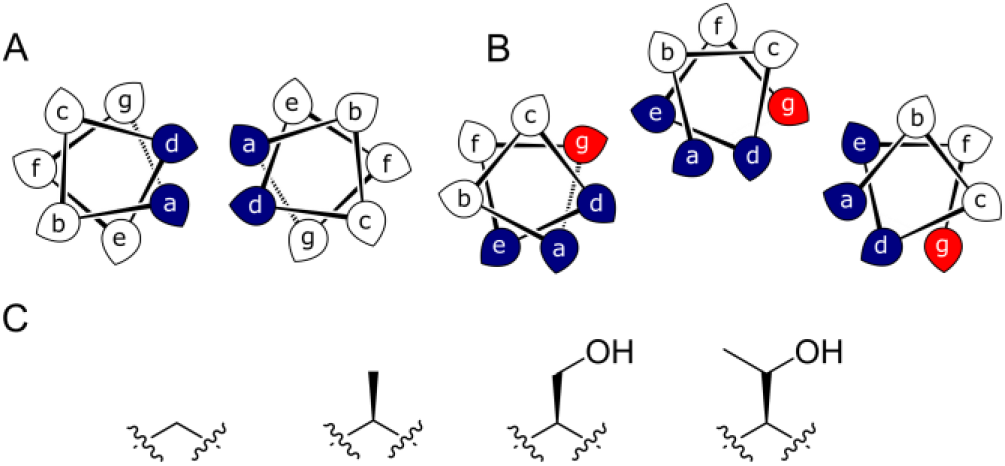
**A&B**: Type-N (**A**) and Type-2 (**B**) CC interfaces. The ***g*** position is highlighted in the Type-2 interface. **C**: Side-chain structures of glycine, alanine, serine and threonine.

**Table 1.**
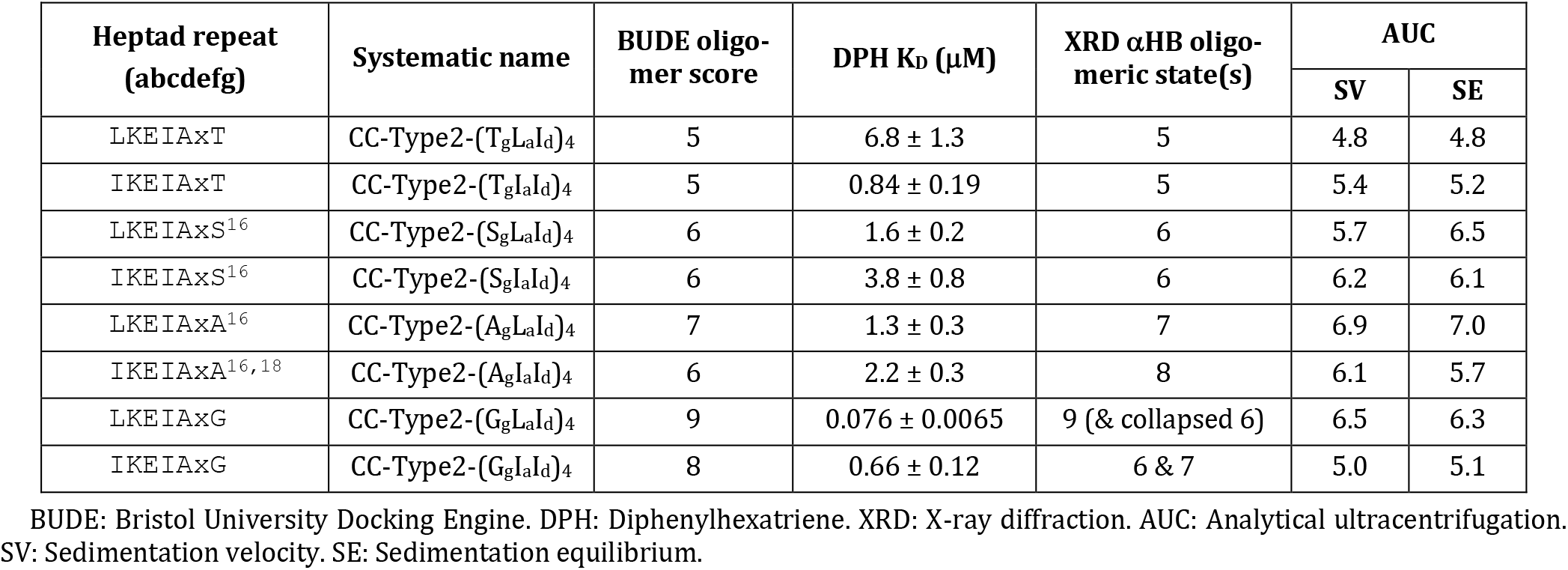
Sequences of *de novo* αHBs and summary of biophysical characterization.

These previously described αHBs show a trend: increasing the size of side chains at ***g*** decreases the oligomer state formed. Therefore, we reasoned that a series of minimal changes, solely at the ***g*** position—*i.e.,* at Z in L/IxxIAxZ repeats—might direct oligomeric state systematically and reliably. Specifically, we considered the addition of a single heavy atom (C or O) to the side chain through the series glycine (Gly, G), Ala, Ser and threonine (Thr, T) (Figure 1C). Our aim was to make a series of peptides with minimal changes to give a robust family of αHBs with tunable oligomeric state for applications in protein design.^13^

To supplement foregoing designs with Ser and Ala at ***g***, we designed four peptides with Gly or Thr at this position; *i.e.,* CC-Type2-(G_g_L_a_I_d_)_4_, CC-Type2-(G_g_I_a_I_d_)_4_, CC-Type2-(T_g_L_a_I_d_)_4_ and CC-Type2-(T_g_I_a_I_d_)_4_, Tables 1 and S1. For simplicity, we refer to these as Gly@***g*** and Thr@***g*** peptides, respectively. Our hypothesis was that these should direct larger and smaller oligomers, respectively.

We built and optimized parametric models for Gly@***g*** and Thr@***g*** in ISAMBARD.^19^ The sequences were modelled as parallel αHBs of oligomer state 5 to 10, and scored using BUDE^20–21^ (Table S3). For the historical designs, the most-favored states were indeed those observed experimentally,^16^ the exception being CC-Type2-(A_g_I_a_I_d_A_e_)_4_, which predicted as a hexamer as observed in solution, but crystallizes as an octamer.^18^ Encouragingly, the new Thr@***g*** sequences consistently scored best as pentamers: for the ***a*** = ***d*** = Ile variant the pentameric assembly was favored outright; while the ***a*** = Leu, ***d*** = Ile variant scored equally well as pentamer or hexamer. Conversely, both Gly variants scored more favorably as larger oligomeric states: the ***a*** = ***d*** = Ile variant as an octamer; and ***a*** = Leu, ***d*** = Ile as a nonamer. Although, there was less discrimination between models for the Gly@***g*** sequences than for Thr@***g***, Figure S1. Thus, modeling supports the hypothesis that oligomeric states of Type-2 CCs can be tuned by side chains at ***g***.

The Gly@***g*** and Thr@***g*** peptides were synthesized, purified by HPLC, and confirmed by mass spectrometry (Figures S2-8).

Circular dichroism (CD) spectroscopy indicated that all four peptides were α helical at low μM concentrations, Figure 2A. CD spectra recorded at increasing temperatures showed that both Thr@***g*** variants were hyperthermostable, Figure 2B. Whereas, the Gly@***g*** variant with ***a*** = Leu, ***d*** = Ile had the beginnings of a thermal unfolding curve consistent with the anticipated destabilizing effect of Gly on α-helical structures.^22–24^ The Gly@***g*** variant with ***a*** = ***d*** = Ile unfolded at 45 °C and did not fully refold on cooling.

**Figure 2.**
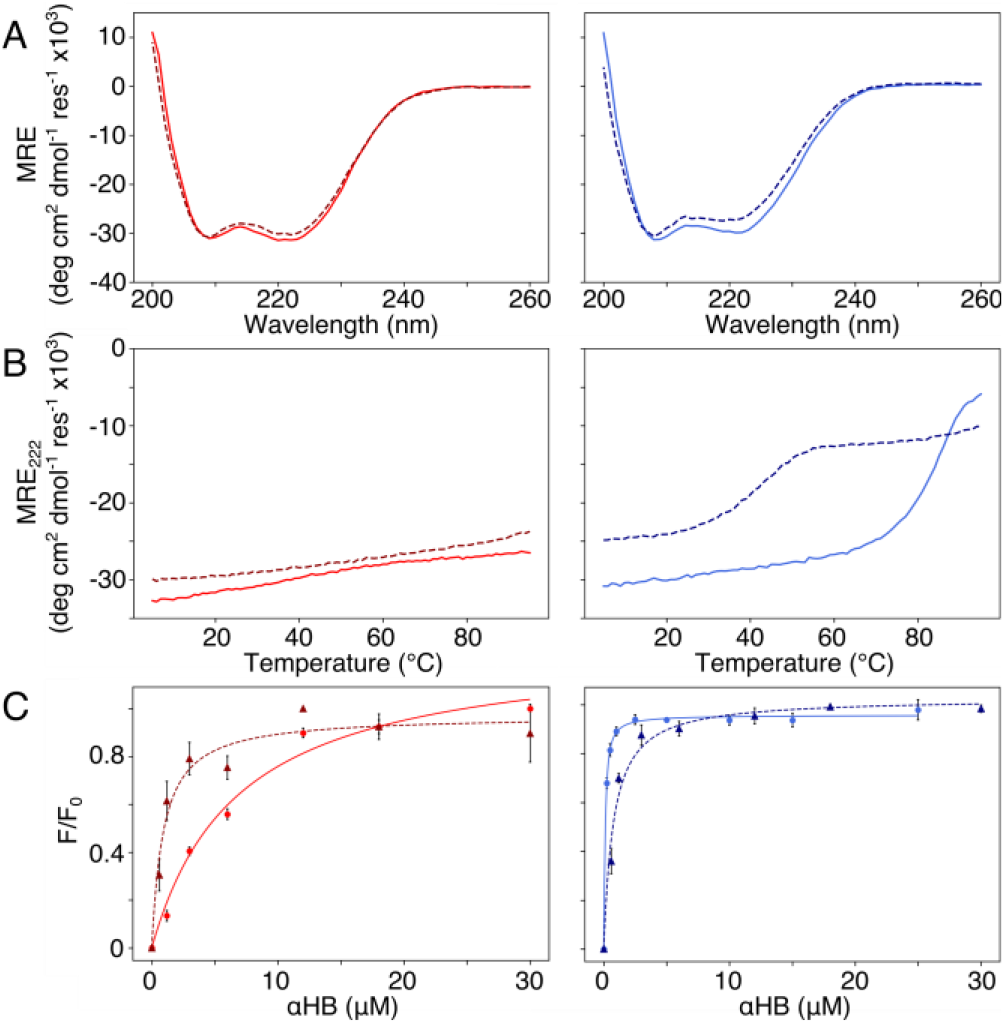
Solution-phase biophysical characterization of Gly@***g*** (right) and Thr@***g*** (left) peptides. **A**: CD spectra at 20 °C. **B:** Thermal denaturation following the CD signal at 222 nm (MRE_222_). **C:** Saturation binding curves with DPH. **Key:** CC-Type2-(T_g_L_a_I_d_)_4_ (red), CC-Type2-(T_g_I_a_I_d_)_4_ (dark red, dashed), CC-Type2-(G_g_L_a_I_d_)_4_ (blue) and CC-Type2-(G_g_I_a_I_d_)_4_ (navy, dashed). **Conditions**: **A & B**, 10 μM peptide. **C**, 0 - 300 μM peptide, 1 μM DPH, 5% v/v DMSO. All experiments were performed in phosphate buffered saline at pH 7.4 (PBS; 8.2 mM Na_2_HPO_4_, 1.8 mM KH_2_PO_4_, 137 mM NaCl, 2.4 mM KCl).

Next, dye-binding assays were used to assess the presence of accessible channels, Figures 2C & S9 and Table 1.^16^, ^18^ All four peptides bound diphenylhexatriene, DPH, consistent with formation of αHBs.

We determined X-ray protein crystal structures for all Gly@,***g*** and Thr@,***g*** variants. As predicted, the Thr@,***g*** peptides formed parallel pentamers (Figure 3 and Tables S4&5). Comparison with the foregoing computationally designed pentamer— CC-Type2-(I_g_L_a_I_d_E_e_)_4_^16^—revealed similar CC parameters for the structures. Significantly, Thr@,***g*** with ***a*** = ***d*** = Ile has a wider pore of ≈9 Å than previous designs (≈7 Å), increasing scope to functionalize this variant.

**Figure 3.**
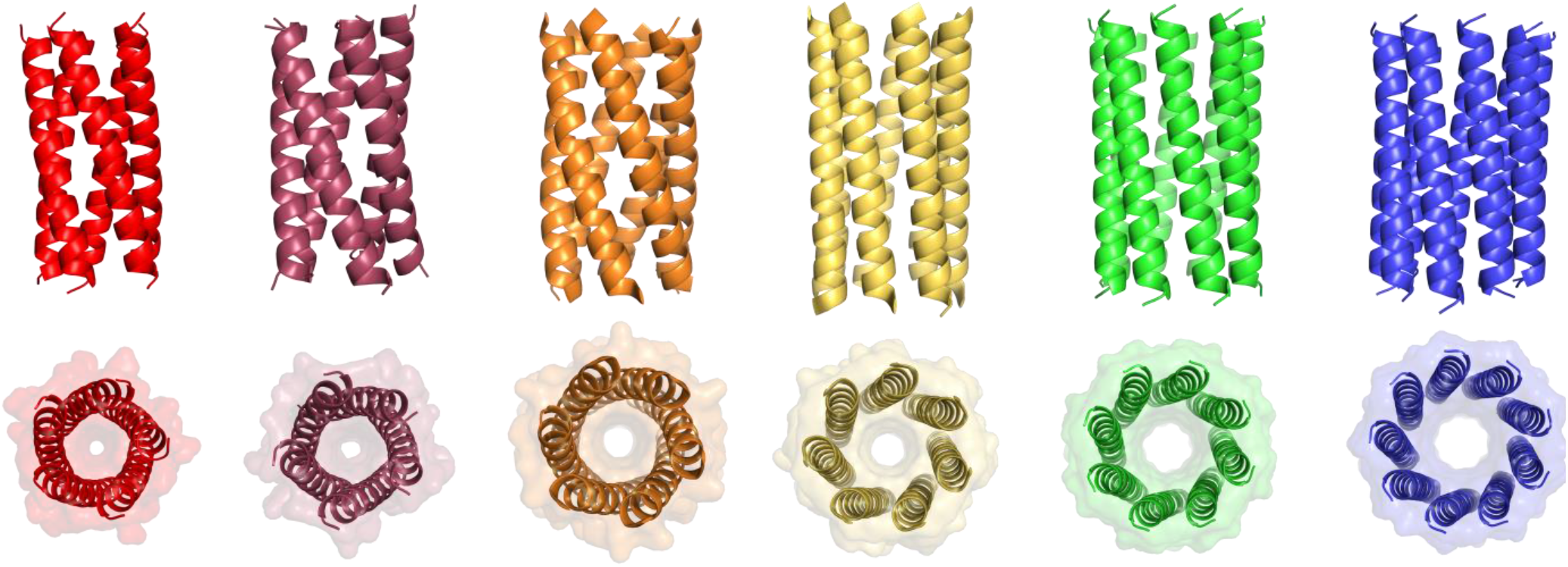
X-ray crystal structures for pentameric through to nonameric αHBs. Left to right: CC-Type2-(T_g_L_a_I_d_)_4_ (red, PDB: 7BAS), CC-Type2-(T_g_I_a_I_d_)_4_ (dark red, PDB: 7BAU), CC-Type2-(S_g_L_a_I_d_)_4_ (orange, PDB: 4PN9), CC-Type2-(A_g_L_a_I_d_)_4_ (yellow, PDB: 4PNA), CC-Type2-(A_g_I_a_I_d_)_4_ (green, PDB: 6G67) and CC-Type2-(G_g_L_a_I_d_)_4_ (blue, PDB: 7BIM).

The Gly@,***g*** peptide with ***a*** = Leu, ***d*** = Ile crystallized in two forms. Gratifyingly, one was an all-parallel nonamer, which is a new αHB with an exceptionally large pore ≈9.5 – 11.5 Å across, Figure 3 (Table S4). Attempts to model solvent into density observed in the channel were inconsistent. Therefore, representative solvent molecules were included where they match the density; though these do not make any stabilizing contacts with protein. Although there are natural nonameric protein assemblies,^25–26^ this is the first stand-alone α-helical CC of this size. The second crystal form revealed a collapsed C_2_-symmetric 6-helix bundle (Figure 4A and B, Table S4). This was surprising, as β-branched residues at ***d*** usually prevent collapse.^18^ We posit that the small size Gly relaxes this design rule allowing access to other parts of the CC free-energy landscape.

**Figure 4.**
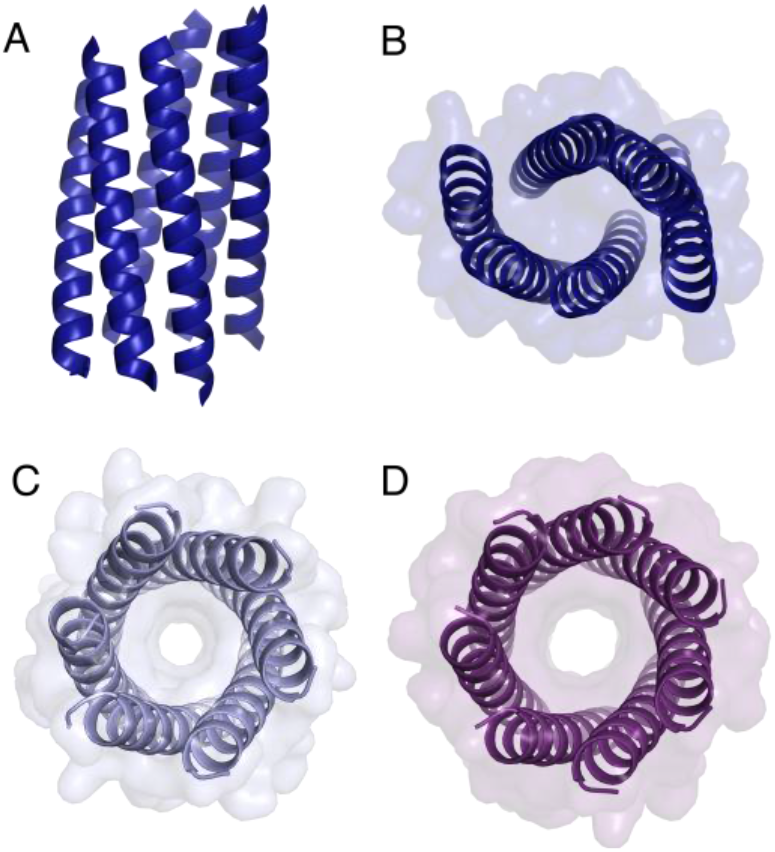
Structures of Gly@,***g*** varaints. **A&B:** Orthogonal views of the collapsed hexameric-form of CC-Type2-(G_g_L_a_I_d_)_4_ (PDB: 7A1T). **C&D**: The hexameric (**C**) and heptameric (**D**) form of CC-Type2-(G_g_I_a_I_d_)_4_ (PDB: 7BAT & 7BAW).

Gly@,***g*** with ***a*** = ***d*** = Ile also crystallized in two forms (Figure 4C and D, Table S5), However, both solved as αHBs; a hexamer and a heptamer. Thus, β-branched Ile at both ***a*** and ***d*** maintains the open assembly^18^ even with Gly residues. Although a larger oligomer was predicted *in silico*, the calculated internal energies for Gly@,***g*** are similar for the different oligomers (Figure S1).

Because of the apparent structural duality with Gly@,***g***, we examined the oligomeric states of all peptides in solution by analytical ultracentrifugation (AUC; Table 1, Figures S10 – S13). Both sedimentation velocity (SV) and sedimentation equilibrium (SE) measurements for the Thr@,***g*** variants returned pentameric weights consistent with the X-ray crystal structures. Indeed, for the pentamers through heptamers for these and previous designs, the correlation between the solution-phase and crystal-state oligomers is good, Table 1. However, where larger oligomers (octamer^18^ and nonamer) are observed in crystals, the correlation is poor with smaller oligomers consistently observed in solution, Table 1. This suggests that the solution states are the dominant species, and that the higher oligomers observed by X-ray crystallography are meta-stable. This is consistent with smaller oligomers being entropically favored. In addition, the crystallization conditions for the octamer and nonamer contained isopropanol. This increases the hydrophobicity of the bulk solvent, which potentially supports larger, hydrophobic pores that would otherwise be energetically unfavorable. Nevertheless, these are legitimate states to consider as they are clearly accessible on the CC free-energy landscape.

Summarizing these data, Type-2 CC peptides with sequence repeats LppIApZ and Z = Thr, Ser, or Ala, form pentameric, hexameric, and heptameric αHBs, respectively. Adding Gly to the series accesses a nonamer, but only in the crystal state. Similarly, an IppIApA peptide forms an octamer in the crystal state.^18^ These X-ray crystal structures enabled us to examine the structural transitions in detail.

In all of the structures, the ***a*** and ***d*** residues contribute both to the lumens and to interactions between neighboring helices. SOCKET^27–28^ analysis revealed that residues at ***a*** form knobs that fit into holes made by ***d’_-1_-g’_-1_-a’-d’*** of a neighboring helix.^29^ These knobs are complemented by interactions formed by the residues at ***g’*** varied herein. The Cα-to-Cβ bond vectors of side chains at ***g*** point directly towards the adjacent helix, Figure 5A - C. In classical CCs, this is called perpendicular packing, and it restricts how close the helices can approach.^3^, ^29^ Thus, mutations at ***g*** might be expected to influence the quaternary structure. The changes made in the Gly→Ala→Ser→Thr series progressively add a single heavy atom to that side chain: Gly (0 atoms)→Ala (1)→Ser (2)→Thr (3), Figure 1C. It is gratifying, but still surprising, that this leads to unitary changes in oligomer state, at least for the Ala, Ser, and Thr variants. This is manifest in adjacent helix-helix distances through the series, Figure 4D. The average distance increases from 8.0 ± 0.1 Å in the Gly*@**g*** nonamer to 10.4 ± 0.1 Å in the Thr*@**g*** pentamer. Thus, through the series, neighboring helices are pushed apart to expel helices from the assembly.

**Figure 5.**
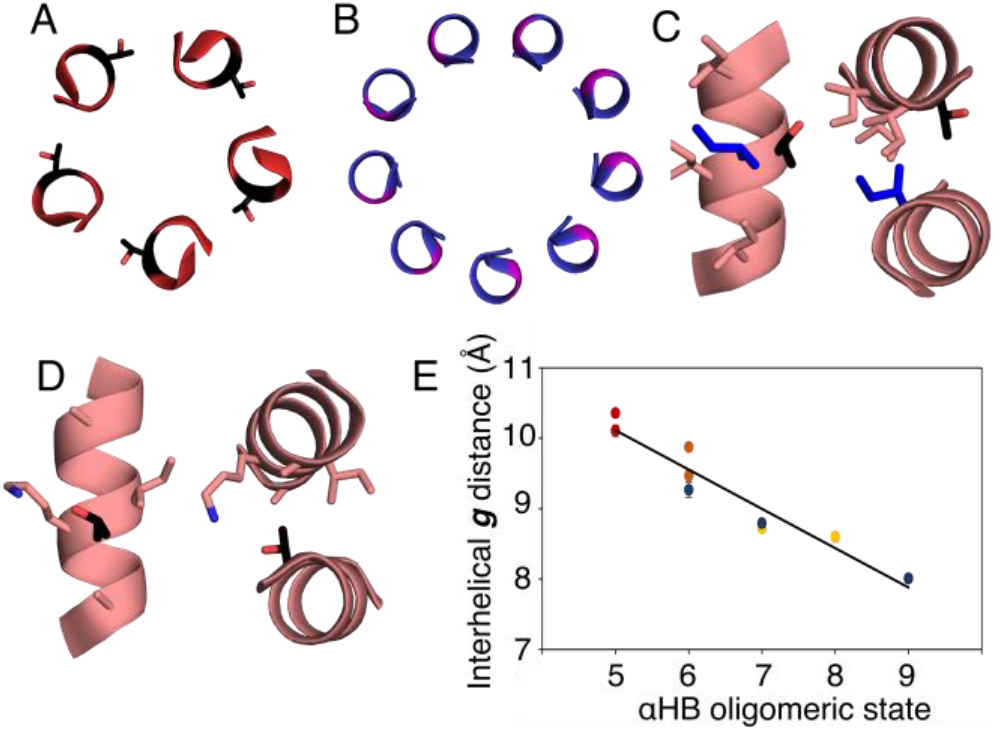
Analysis of pentameric to nonameric αHBs. **A&B**: Cross-sections through CC-Type2-(T_g_I_a_I_d_)_4_ (red) and CC-Type2-(G_g_L_a_I_d_)_4_ (blue) showing the geometry of residues at ***g*** (black and pink, respectively). **C&D**: Knobs-into-holes interaction showing how these residues (black) contribute to a ***d’_-1_-g’_-1_-a’-d’*** hole when Ile is the knob residue (blue) (**C**); and when the Thr at the ***g*** position is a knob residue (**D**). **E**: Interhelical distances in the αHBs with Thr (red), Ser (orange), Ala (yellow) and Gly (blue) at ***g***. Errors are the standard deviation of measurements from the central heptads of each structure.

The judicious placement of Gly may prove useful in designing αHBs to unlock previously unseen architectures. However, using Gly presents challenges that must be met to allow its full exploitation. The first challenge is incorporating multistate design into αHBs, *i.e.* considering multiple conformations and/or assemblies that may become accessible in the CC free-energy landscape. This approach is being applied to other systems.^30–32^ It is tractable to model many possible Type-2 αHBs to direct computational design.^16^ However, this becomes difficult with increasing off-target states, *e.g.* collapsed and anti-parallel structures. The second challenge is to stabilize the larger, and clearly accessible, oligomer states in solution. One possibility would be to introduce networks of polar residues to reduce the penalty of all-hydrophobic channels. *De novo* αHBs have proved viable candidates for functional protein design.^11–13^ Reliably accessing scaffolds with significantly larger pores systematically and with minimal changes in primary sequence, would expand the scope for this and future applications.

In conclusion, we have combined rational design, computational modelling, and structural biology for a series of αHBs with mutations from Gly→Ala→Ser→Thr at all of the ***g*** sites in a CC sequence. Minimal and stepwise changes in size of the residue at these sites, combined with Leu/Ile at ***a*** and Ile at ***d***, control the oligomeric state of the assembly. This expands the range of αHBs that can be designed systematically from pentamer (with Thr at ***g***) to a nonamer (with Gly at ***g***). Inspection of X-ray crystal structures rationalizes the role of side-chain bulk at ***g*** in dictating inter-helical packing distance, angles, and, thus, oligomeric state. Gly***@*g** is the first example of a stand-alone nonameric CC. However, it appears that such high oligomeric states (8 and 9) are on the edge of what is possible for Type-2 CC sequences as they are not favored in solution.^16^, ^18^, ^33–35^ Nonetheless, the X-ray crystal structures show that they are accessible. The rarity of such assemblies in nature^6^, ^17^, ^25–26^ and their potential as scaffolds for functional *de novo* design^8–11^ makes these large αHBs tantalizing targets for design.

## Notes

X-ray crystal structures for CC-Type2-(T_g_L_a_I_d_)_4_-W19BrPhe, CC-Type2-(T_g_I_a_I_d_)_4_-W19BrPhe, CC-Type2-(G_g_L_a_I_d_)_4_, CC-Type2-(G_g_L_a_I_d_)_4_-W19BrPhe and CC-Type2-(G_g_I_a_I_d_)_4_ are available from the Protein Data Bank. Accession codes: 7AIT, 7BAS, 7BAT, 7BAU, 7BAV, 7BAW & 7BIM.

## ACKNOWLEDGMENT

WMD, GGR and DNW were supported by a European Research Council Advanced Grant (340764). FOJM is supported by the Bristol Chemical Synthesis Centre for Doctoral Training funded through the EPSRC (EP/G036764). KLS is supported by the South West Biosciences Doctoral Training Partnership through the BBSRC (BB/M009122/1). DNW is supported by a BBSRC responsive-mode grant (BB/R00661X/1). We thank the University of Bristol School of Chemistry Mass Spectrometry Facility for access to the EPSRC-funded Bruker Ultraflex MALDI-TOF instrument (EP/K03927X/1) and BrisSynBio for access to the BBSRC-funded BMG Labtech Clariostar Plate Reader (BB/L01386X/1). We would like to thank Diamond Light Source for access to beamlines I03, I04, I04-1 and I24 (Proposal 12342 & 23269), and for the support from the macromolecular crystallography staff.

## Notes

### Competing Interest Statement

The authors have declared no competing interest.

