## Supplementary Data for "Coiled coils 9-to-5: Rational *de novo* design of α-helical barrels with tunable oligomeric states"

### 1 Supplementary Methods

#### 1.1 General

Solid-phase peptide synthesis (SPPS) reagents were purchased from Cambridge Reagents with the exception of N,N'-diisopropylcarbodiimide (DIC) purchased from Carbosynth and Rink amide MBHA resin from Merck. All other chemicals were purchased from Merck. All biophysical experiments were conducted in phosphate buffered saline (PBS; 10 mM sodium phosphate dibasic, 1.8 mM potassium phosphate dibasic, 137 mM sodium chloride, 2.4 mM potassium chloride, pH 7.4). Peptide concentration was determined at 280 nm using a Nanodrop 2000 (ThermoScientific) spectrometer ($\varepsilon_{280}$ = 5690 $cm^{-1}$) or at 214 nm using a Cary-100 (Agilent) UV-Vis spectrometer by measuring the peptide bond.[1] Peptide characterisation data for CC-Type2-($S_gL_aI_d$)$_4$, CC-Type2-($S_gI_aI_d$)$_4$, CC-Type2-($A_gL_aI_d$)$_4$ and CC-Type2-($A_gI_aI_d$)$_4$ has been published previously.[2-3]

#### 1.2 Oligomer state prediction

Parametric models were built using the coiled coil classification in ISAMBARD.[4] Each sequence (Table S1) was optimised and scored as an $\alpha$HB with oligomer state between 5 and 10 inclusive using the in-build GA optimizer and the BUFF force-field.

#### 1.3 Solid-phase peptide synthesis

SPPS was performed on a Liberty Blue automated peptide synthesizer (CEM) with inline UV monitoring. All peptides were synthesized as the C-terminal amide on Rink amide MBHA resin using standard Fmoc-chemistry, with DIC/6-chloro-1-hydroxybenzotriazole as the coupling reagents. Fmoc was removed using 20% v/v morpholine:dimethylformamide (DMF). All peptides were acetyl capped through addition of pyridine (0.5 mL) and acetic anhydride (0.3 mL) in DMF (9.2 mL), shaking at room temperature (rt) for 20 minutes. Peptides were cleaved from the resin with addition of 95:2.5:2.5 v/v trifluoroacetic acid (TFA):$H_2O$:triisopropylsilane, shaking at rt for 3 hours before removal by $N_2$ blow down. Cleaved peptide was precipitated with cold diethyl ether, isolated via centrifugation and dissolved in 50:50 v/v acetonitrile (MeCN):$H_2O$. Crude peptides were lyophilized to yield a white or off-white powder.

#### 1.4 Peptide purification and characterisation

All peptides were purified by reverse phase HPLC (JASCO) using a Luna C18 (Phenomenex) column (150 x 10 mm, 5 $\mu$M particle size, 100 Å pore size). Crude peptide was dissolved at 5 mg/mL in 40% v/v MeCN in $H_2O$ with 0.1% TFA. A 40-100% gradient of MeCN in $H_2O$ with 0.1% TFA over 30 minutes was used to separate the target peptide before confirmation by analytical HPLC and MALDI-TOF.

Analytical HPLC was performed as above using a Kinetix C18 (Phenomenex) column (100 x 4.6 mm, 5 μM particle size, 100 Å pore size) over a 25 minute gradient.

MALDI-TOF was performed on an Ultraflex MALDI-TOF (Bruker) in positive-ion reflector mode. Samples were spotted on a ground steel plate using dihydroxybenzoic acid or α-cyan-4-hydroxycinnamic. Masses quoted are for the monoisotopic mass of the singly protonated species $[M+H^+]$.

#### 1.5 Circular dichroism spectroscopy

Circular dichroism (CD) spectroscopy was performed on a Jasco J-810 or J-815 spectropolarimeter fitted with a Peltier temperature controller. Data were collected in a 5 mm quartz cuvette between 190 and 260 nm (100 nm $min^{-1}$, 1 nm interval and bandwidth, 1s response time). CD spectra were acquired at 10 μM peptide concentration at 20 °C. Thermal denaturation spectra were collected at 222 nm using the settings and peptide concentration as above between 5 and 95 °C.

#### 1.6 Analytical ultracentrifugation

Analytical ultracentrifugation (AUC) was performed on a Beckman Optima X-LA or X-LI analytical ultracentrifuge with an An-50-Ti or An-60-Ti rotor (Beckman-Coulter). Buffer densities, viscosities and peptide partial specific volumes ($\bar{v}$) were calculated using SEDNTERP (http://rasmb.org/sednterp/).

Sedimentation equilibrium experiments were performed at 70 μM peptide concentration at 20 °C in 6-channel epon centrepieces with quartz windows. Data were collected between 15-30 krpm with a minimum of 3 speeds sampled after equilibration for 8 hours. Data were fitted using SEDPHAT to a single species model.[5] Monte Carlo analysis was performed to give 95% confidence limits.

Sedimentation velocity (SV) experiments were performed at 150 μM peptide concentration at 20 °C in 2-channel epon or aluminium centrepieces with quartz windows. Data were collected at 50 krpm at 5-minute intervals for a total of 120 scans and fitted to a continuous distribution using SEDFIT[2] at a 95% confidence level. Residuals are shown as a bitmap with scans ordered vertically from the top of the image. Greyscale shade indicates difference between the model and raw data over the radial range of the fit (residuals <-0.05 black, > 0.05 white).

#### 1.7 Ligand binding

Ligand binding experiments were performed on an epMotion 5070 liquid handler (Eppendorf). The total concentration of ligand was kept constant (0.1 μM in 5% v/v DMSO) and the concentration of αHB varied from 0 – 50 μM. Data were collected on a Clariostar plate reader

(BMG Labtech) using an excitation wavelength of 350 nm and emission monitored at 450 nm. Binding constants were extracted by fitting Equation 1 or Equation 2[6] to the data in SigmaPlot 13.0.

$$y = \frac{B_{max} \cdot x}{K_D + x} \qquad \text{Equation 1}$$

$$y = B_{max} \frac{(c + x + K_D) + \sqrt{(c + x + K_D)^2 - 4cx}}{2c} \qquad \text{Equation 2}$$

Where c is the total concentration of the constant component (*e.g.* DPH), x is the concentration of variable component (*e.g.* $\alpha$HB), $B_{max}$ is the fluorescence signal when all of the constant component is bound, and y is the fraction of bound component being monitored via fluorescence signal.

#### 1.8 X-ray crystal structure determination

Freeze-dried peptides were resuspended in deionised water to approximate concentrations of 10 mg $ml^{-1}$ or 5 mg $ml^{-1}$ for CC-Type2-($T_gL_aI_d$)$_4$-W19BrPhe and CC-Type2-($T_gI_aI_d$)$_4$-W19BrPhe, for vapour-diffusion crystallisation trials using standard commercial screens (JCSG-plus™, Structure Screen 1 + 2, ProPlex™, Morpheus® and PACT Premier™) at 19 °C with 0.3 µl of the peptide solution equilibrated with 0.3 µl of the screen solution. Final crystallisation conditions for all peptides are provided in Table S1. To aid with cryoprotection, crystals were soaked in their respective reservoir solutions containing 25% glycerol prior to freezing. X-ray diffraction data were collected at the Diamond Light Source (Didcot, UK) on beamline I04, at various wavelengths. Data were processed using automated or manual pipelines. Automated pipelines: Xia2 pipelines,[7] which ports data through DIALS[8] or MOSFLM[9] to POINTLESS and AIMLESS[10] as implemented in the CCP4 suite,[11] or XDS to XSCALE;[12] or the AutoPROC pipelines, which use the same integrating and data reduction software in addition to STARANISO.[13] Manually: The images for CC-Type2-($T_gI_aI_d$) were manually reprocessed using the Dials User Interface[14] and run through POINTLESS and AIMLESS in the CCP4 suite. CC-Type2-($G_gL_aI_d$)$_4$ collapsed hexamer was experimentally phased using Br-atom SAD phasing using the automated Big EP pipeline that experimentally phases and builds using AUTOSHARP,[15] AUTOSOL[16] and Crank2.[17] CC-Type2-($T_gI_aI_d$), CC-Type2-($G_gL_aI_d$)$_4$ nonamer and CC-Type2-($G_gI_aI_d$) hexamer were phased using *ab initio* phasing using ARCIMBOLDO_LITE.[18] CC-Type2-($G_gI_aI_d$) heptamer, was phased using FRAGON,[19] the initial phases were modelled into and refined using ARP/wARP.[20] All other structures were solved by molecular replacement using full or partial poly-alanine models (as dictated by the Matthews Coefficient), generated from existing coiled-coil structures, using PHASER.[21] Final structures were obtained after iterative rounds of model building with COOT[22] and refinement with REFMAC5.[23] Late-stage models of all structures were submitted to PDB_REDO[24] and further refined with REFMAC5. Solvent-exposed atoms lacking map density were either deleted or left at full occupancy. Data collection and refinement statistics are provided in Tables S4 and S5.

**Table S1** Crystallisation conditions used to obtain the structures discussed in this article.

| Sequence | Crystallisation conditions* | Molecular dimensions screen |
|---|---|---|
| **CC-Type2-($T_gL_aI_d$)$_4$-W19BrPhe** (7BAS) | 50 mM MES buffer, 5 % w/v PEG 5000 MME, and 6 % v/v 1-propanol at pH 6.5 | Proplex D2 |
| **CC-Type2-($T_gL_aI_d$)$_4$-W19BrPhe** (7BAV) | 50 mM potassium chloride 50 mM sodium HEPES buffer, and 7.5 % w/v PEG 5000 MME at pH 7.0 | Proplex D3 |
| **CC-Type2-($T_gI_aI_d$)$_4$ W19BrPhe** (7BAU) | 1.0 M Ammonium formate, 50 mM sodium HEPES buffer at pH 7.5 | Structure screen 1 and 2 F3 |
| **CC-Type2-($G_gL_aI_d$)$_4$ (nonameric αHB)** (7BIM) | 50 mM sodium HEPES buffer, 100 mM ammonium acetate and 12.5% (v/v) isopropyl alcohol at pH 7.5 | Proplex H11 |
| **CC-Type2-($G_gL_aI_d$)$_4$ (collapsed hexameric bundle)** (7AIT) | 50 mM sodium HEPES buffer, 500 mM sodium acetate and 25 mM cadmium sulfate (8/3)-hydrate at pH 7.5 | Structure screen 1 and 2 F4 |
| **CC-Type2-($G_gI_aI_d$)$_4$ (hexameric αHB)** (7BAT) | 50 mM MES buffer, 5 % w/v PEG 5000 monomethyl ether, and 6 % v/v 1-propanol at pH 6.5 | Proplex D2 |
| **CC-Type2-($G_gI_aI_d$)$_4$ (heptameric αHB)** (7BAW) | 25 mM imidazole, 25 mM MES monohydrate (acid) buffer, 15 mM sodium nitrate, 15 mM sodium phosphate dibasic, 15 mM ammonium sulfate, 10% v/v ethylene glycol, and 5 % w/v PEG 8000 at pH 6.5 | Morpheus C2 |

*These are the dispensed concentrations based on a 1:1 dilution with the peptide solution.

#### 2 Supplementary Data

**Table S2.** Sequences of *de novo* peptides in this study

| **Systematic Name** | **Heptad repeat** (*gabcdef*) | **Alternative Name** | **Sequence** |
|---|---|---|---|
| CC-Type2-($T_gL_aI_d$)$_4$ | TLKEIAx | - | Ac-GEIAQTLKEIAKTLKEIAWTLKEIAQTLKG-NH$_2$ |
| CC-Type2-($T_gL_aI_d$)$_4$-W19BrPhe | TLKEIAx | - | Ac-GEIAQTLKEIAKTLKEIAΦTLKEIAQTLKG-NH$_2$ |
| CC-Type2-($T_gI_aI_d$)$_4$ | TIKEIAx | - | Ac-GEIAQTIKEIAKTIKEIAWTIKEIAQTIKG-NH$_2$ |
| CC-Type2-($T_gI_aI_d$)$_4$-W19BrPhe | TIKEIAx | - | Ac-GEIAQTIKEIAKTIKEIAΦTIKEIAQTIKG-NH$_2$ |
| CC-Type2-($S_gL_aI_d$)$_4$ | SLKEIAx | CCHex2 | Ac-GEIAKSLKEIAKSLKEIAWSLKEIAKSLKG-NH$_2$ |
| CC-Type2-($S_gI_aI_d$)$_4$ | SIKEIAx | CCHex3 | Ac-GEIAQSIKEIAKSIKEIAWSIKEIAQSIKG-NH$_2$ |
| CC-Type2-($A_gL_aI_d$)$_4$ | ALKEIAx | CCHept | Ac-GEIAQALKEIAKALKEIAWALKEIAQALKG-NH$_2$ |
| CC-Type2-($A_gI_aI_d$)$_4$ | AIKEIAx | - | Ac-GEIAQAIKEIAKAIKEIAWAIKEIAQAIKG-NH$_2$ |
| CC-Type2-($G_gL_aI_d$)$_4$ | GLKEIAx | - | Ac-GEIAQGLKEIAKGLKEIAWGLKEIAQGLKG-NH$_2$ |
| CC-Type2-($G_gI_aI_d$)$_4$ | GLKEIAx | - | Ac-GEIAQGIKEIAKGIKEIAWGIKEIAQGIKG-NH$_2$ |
| CC-Type2-($G_gL_aI_d$)$_4$-W19BrPhe | GLKEIAx | - | Ac-GEIAQGLKEIAKGLKEIAΦGLKEIAQGLKG-NH$_2$ |

Φ = 4-bromo-phenylalanine

**Table S3.** *In silico* modelling predicted scores and parameters

| Heptad repeat | Oligomer state | BUDE score | Next oligomer | BUDE score difference | Parameters of best scoring model | | |
|---|---|---|---|---|---|---|---|
| | | | | | Phi-Cα (°) | Radius (Å) | Pitch (Å) |
| TLKEIAx | 6* | -554.8 ± 1.4 | 5* | 0 | 219.4 | 9.1 | 211.6 |
| TIKEIAx | 5 | -562.1 + 0.0 | 6 | -14.1 | 217.5 | 8.1 | 120.4 |
| GLKEIAx | 9 | -565.7 ± 0.2 | 8 | -3.1 | 218.6 | 11.4 | 261.23 |
| GIKEIAx | 8 | 574.2 ± 0.0 | 10 | -3.8 | 222.0 | 10.6 | 150.0 |

*TLKEIAx scored equally well as a pentamer or hexamer. The best scoring individual model was hexameric.

**Table S4. Merging and refinement statistics for all X-ray crystal structures.**

| | **CC-Type2-($T_gL_aI_d$)$_4$-W19BrPhe** | **CC-Type2-($T_gL_aI_d$)$_4$-W19BrPhe** | **CC-Type2-($T_gI_aI_d$)$_4$-W19BrPhe** | **CC-Type2-($G_gL_aI_d$)$_4$ (nonameric aHB)** |
|---|---|---|---|---|
| PDB Code | 7BAS | 7BAV | 7BAU | 7BIM |
| Wavelength (Å) | 0.8610 | 0.9999 | 0.92 | 0.9795 |
| Resolution range (Å) | 39.62 – 1.1 (1.139 – 1.1) | 37.73 – 1.3 (1.347 – 1.3) | 40.63 – 1.42 (1.45 – 1.42) | 66.85 – 1.64 (1.67 – 1.64) |
| Space group | P 2 21 21 | P 21 21 21 | P 1 21 1 | P 1 21 1 |
| Unit cell lengths (Å) | 43.56 54.65 57.53 | 43.73 51.83 55.04 | 40.60 53.45 56.16 | 71.21 128.08 71.45 |
| Unit cell angles (°) | 90 90 90 | 90 90 90 | 90 90 90 | 90 110.67 90 |
| Total reflections | 1370194 (120688) | 750174 (65493) | 475369 (22901) | 996133 (50464) |
| Unique reflections | 56469 (5564) | 31406 (3051) | 23553 (1132) | 146091 (7322) |
| Multiplicity | 24.3 (21.7) | 23.9 (21.5) | 20.2 (20.2) | 6.8 (6.9) |
| Completeness (%) | 99.16 (99.23) | 99.91 (99.32) | 99.5 (97.3) | 99.9 (100) |
| Mean I/sigma(I) | 6.07 (0.29) | 4.60 (0.20) | 11.0 (1.6) | 14.2 (1.4) |
| Wilson B-factor (Å$^2$) | 9.23 | 15.76 | 13.3 | 18.486 |
| R-merge(I) | 0.246 (6.178) | 0.3253 (63.53) | 0.151 (3.399) | 0.074 (1.429) |
| R-meas(I) | 0.2513 (6.325) | 0.3325 (65.05) | 0.157 (3.485) | 0.08 (1.545) |
| R-pim | 0.05069 (1.344) | 0.06767 (13.84) | 0.048 (1.071) | 0.031 (0.583) |
| CC1/2 | 0.999 (0.398) | 0.998 (0.372) | 0.996 (0.567) | 0.999 (0.526) |
| CC* | 1 | 1 | | |
| Reflections used in refinement | 55998 | 31398 | 42429 | 138782 |
| Reflections used for R-free | 2869 | 1531 | 2341 | 7270 |
| R-work | 0.1915 | 0.2006 | 0.177 | 0.1823 |
| R-free | 0.2212 | 0.2296 | 0.226 | 0.2055 |
| CC(work) | 0.954 | 0.954 | 0.9228 | |
| CC(free) | 0.960 | 0.917 | 0.9009 | |
| Number of non-hydrogen atoms | 1341 | 1285 | 2460 | 8789 |
| macromolecules | 1150 | 1152 | 2288 | 8149 |
| Ligands | 6 | 11 | 132 | 262 |
| Solvent | 185 | 122 | 184 | 378 |
| Protein residues | 150 | 150 | 150 | 1107 |
| RMS(bonds) | 0.018 | 0.018 | 0.0119 | 0.0145 |
| RMS(angles) | 2.01 | 2.16 | 1.555 | 1.6222 |
| Ramachandran favoured (%) | 100 | 100 | 100 | 100 |
| Ramachandran allowed (%) | 0 | 0 | 0 | 0 |
| Ramachandran outliers (%) | 0 | 0 | 0 | 0 |
| Rotamer outliers (%) | 0 | 0 | 0 | 3 / 737 |
| Clashscore | 1.24 | 4.48 | 2 | 3.74 |
| Average B-factor | 15.49 | 24.44 | 20.36 | 33.21 |
| macromolecules | 14.17 | 23.55 | 19.68 | 31.98 |
| ligands | 21.63 | 46.3 | - | 58.78 |
| solvent | 23.55 | 30.87 | 28.78 | 42.17 |
| Number of TLS groups | 5 | 5 | 10 | 36 |

**Table S5. Merging and refinement statistics for all X-ray crystal structures.**

| | **CC-Type2-($G_gL_aI_d$)$_4$ (collapsed hexameric bundle)** | **CC-Type2-($G_gI_aI_d$)$_4$ (hexameric αHB)** | **CC-Type2-($G_gI_aI_d$)$_4$ (heptameric αHB)** |
|---|---|---|---|
| PDB Code | 7A1T | 7BAT | 7BAW |
| Wavelength (Å) | 0.9187 | 0.9795 | 0.9795 |
| Resolution range (Å) | 59.12-1.42 [59.12-6.35] (1.46-1.420) | 40.51 – 1.77 (1.81 – 1.77) | 50.43 – 1.7 (1.771 – 1.71) |
| Space group | C 2 2 21 | I 2 2 2 | P 21 2 21 |
| Unit cell lengths (Å) | 50.64 129.69 59.12 | 39.89 42.74 126.20 | 37.45 50.43 126.61 |
| Unit cell angles (°) | 90 90 90 | 90 90 90 | 90 90 90 |
| Total reflections | 476205 [5443] (35142) | 137174 | 349995 (35241) |
| Unique reflections | 37216 [482] (2721) | 10914 | 26748 (2600) |
| Multiplicity | 12.8 [11.3] (12.9) | 12.6 | 13.1 (13.5) |
| Completeness (%) | 100 [99.7] (100) | 100.0 (99.0) | 99.59 (99.42) |
| Mean I/sigma(I) | 15 [46.9] (1.1) | 21.6 | 22.24 (0.55) |
| Wilson B-factor Å$^2$) | 18.19 | 42.0 | 36.9 |
| R-merge(I) | 0.082 [0.063] (2.470) | 0.04 (3.324) | 0.04571 (2.526) |
| R-meas(I) | 0.085 [0.067] (2.572) | 0.041 (3.474) | 0.04764 (2.623) |
| R-pim | 0.024 [0.020] (0.709) | 0.012 (0.99) | 0.01319 (0.7015) |
| CC1/2 | 0.999 [0.997] (0.502) | 1.000 (0.387) | 1 (0.63) |
| CC* | 1 | | 1 (0.879) |
| Reflections used in refinement | 35293 | 10911 | 26653 (2590) |
| Reflections used for R-free | 1885 | 514 | 1359 (144) |
| R-work | 0.157 | 0.258 | 0.2445 (0.5066) |
| R-free | 0.198 | 0.265 | 0.2727 (0.4964) |
| CC(work) | 0.975 | 0.9344 | 0.943 (0.737) |
| CC(free) | 0.965 | 0.9399 | 0.929 (0.709) |
| Number of non-hydrogen atoms | 1584 | 614 | 1518 |
| macromolecules | 1486 | 584 | 1452 |
| Ligands | 19 | 12 | 51 |
| Solvent | 79 | 21 | 15 |
| Protein residues | 185 | 86 | 208 |
| RMS(bonds) | 0.0214 | 0.0146 | 0.017 |
| RMS(angles) | 2.099 | 1.60 | 1.83 |
| Ramachandran favoured (%) | 100 | 100 | 100 |
| Ramachandran allowed (%) | 0 | 0 | 0 |
| Ramachandran outliers (%) | 0 | 0 | 0 |
| Rotamer outliers (%) | 0 | 0 | 0 |
| Clashscore | 0 | 6 | 3.23 |
| Average B-factor | 28.18 | 52.29 | 44.3 |
| macromolecules | 29.74 | 51.74 | 43.3 |
| ligands | 18.00 | 51.88 | 70.18 |
| solvent | 47.66 | 68.21 | 53.69 |
| Number of TLS groups | 0 | 3 | 7 |

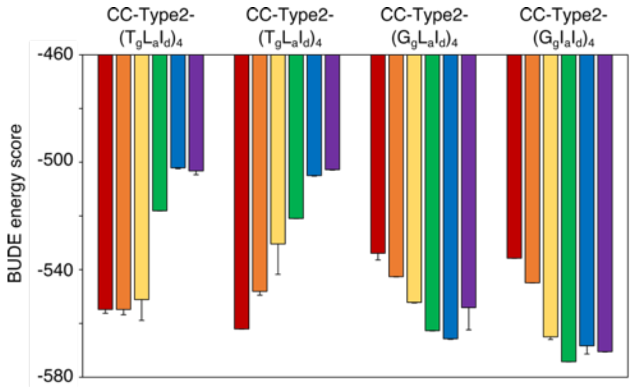

**Figure S1. Rational computational design of αHBs.** BUDE energy scores for each αHB model with 5 – 10 α-helices inclusive is shown (negative energy scores mean a more favourable structure): pentamer (red), hexamer (orange), heptamer (yellow), octamer (green), nonamer (blue) and decamer (purple).

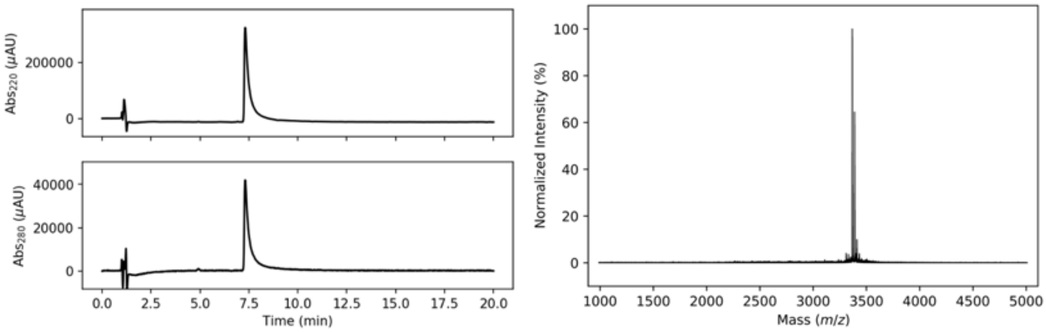

**Figure S2. Characterisation of CC-Type2-($T_gL_aI_d$)$_4$. Left:** Analytical HPLC chromatogram at 220 nm (top) and 280 nm (bottom). **Right:** MALDI-TOF MS. Calculated mass: 3365.93 Da $[M+H]^+$. Observed mass: 3366.05 Da $[M+H]^+$.

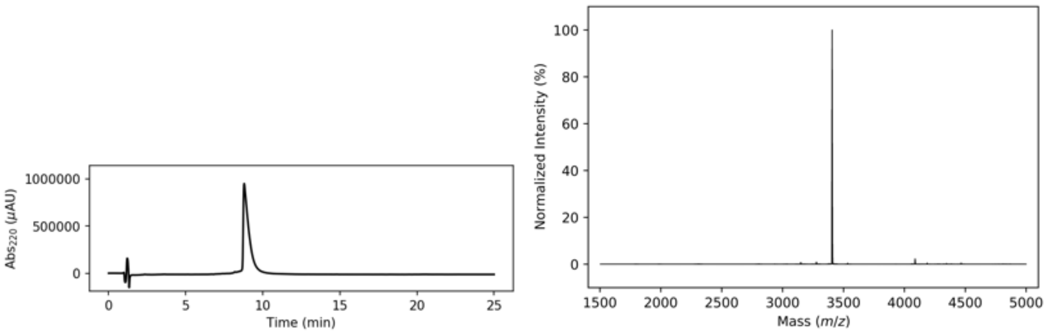

**Figure S3. Characterisation of CC-Type2-($T_gL_aI_d$)$_4$-W19BrPhe. Left:** Analytical HPLC chromatogram at 220 nm (top) and 280 nm (bottom). **Right:** Deconvoluted ESI MS. Calculated mass: 3404.83 Da $[M+H]^+$. Observed mass: 3404.02 Da $[M+H]^+$.

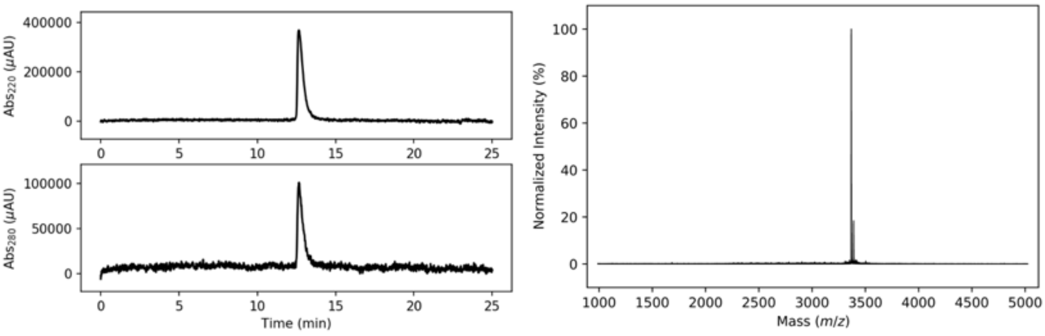

**Figure S4. Characterisation of CC-Type2-($T_gI_aI_d$)$_4$. Left:** Analytical HPLC chromatogram at 220 nm. **Right:** MALDI-TOF MS. Calculated mass: 3365.93 $[M+H]^+$. Observed mass: 3365.916 $[M+H]^+$.

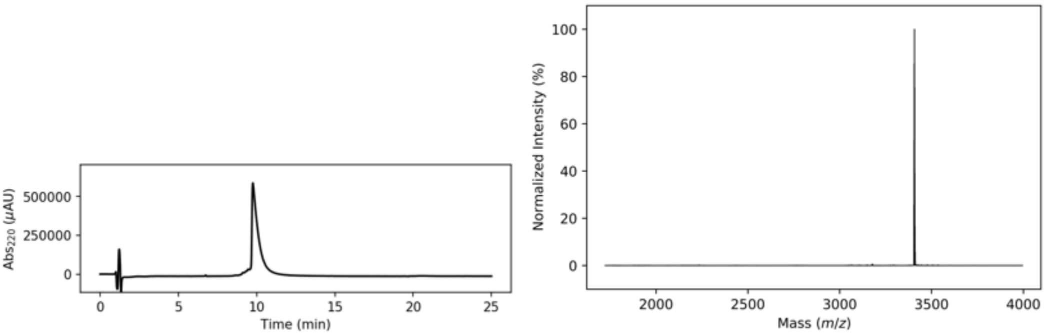

**Figure S5. Characterisation of CC-Type2-($T_gI_aI_d$)$_4$-W19BrPhe. Left:** Analytical HPLC chromatogram at 220 nm. **Right:** Deconvoluted ESI MS. Calculated mass: 3404.83 $[M+H]^+$. Observed mass: 3403.97 $[M+H]^+$.

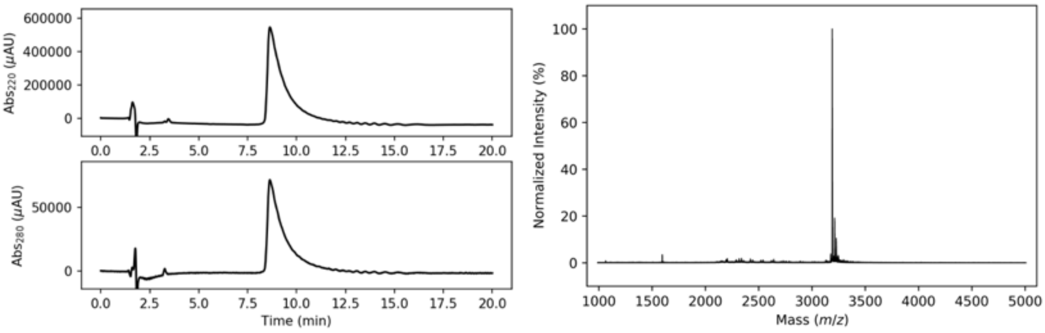

**Figure S6. Characterisation of CC-Type2-($G_gL_aI_d$)$_4$. Left:** Analytical HPLC chromatogram at 220 nm (top) and 280 nm (bottom). **Right:** MALDI-TOF MS. Calculated mass: 3189.825 Da $[M+H]^+$. Observed mass: 3188.853 Da $[M+H]^+$.

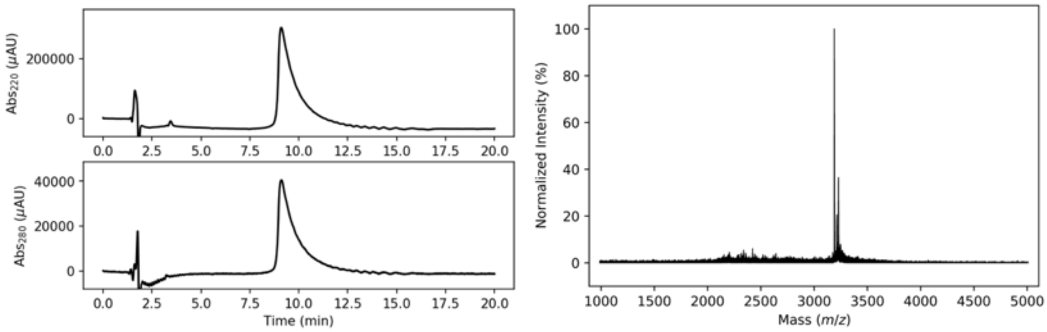

**Figure S7. Characterisation of CC-Type2-($G_gI_aI_d$)$_4$. Left:** Analytical HPLC chromatogram at 220 nm (top) and 280 nm (bottom). **Right:** MALDI-TOF MS. Calculated mass: 3189.825 Da $[M+H]^+$. Observed mass: 3189.788 Da $[M+H]^+$.

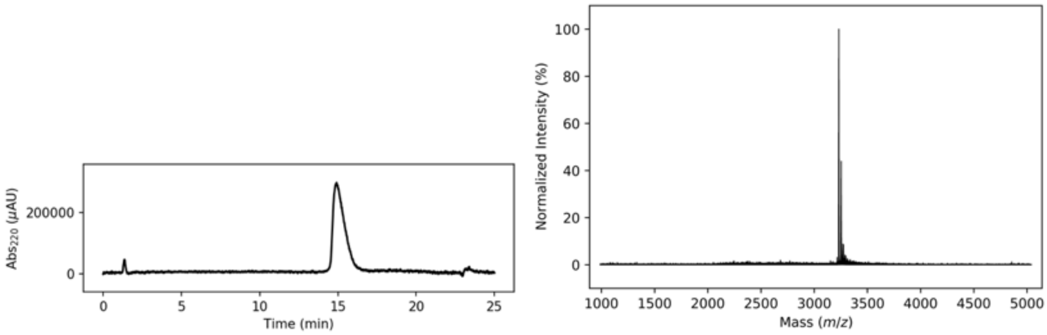

**Figure S8. Characterisation of CC-Type2-($G_gL_aI_d$)$_4$-W19BrPhe. Left:** Analytical HPLC chromatogram at 220 nm (top) and 280 nm (bottom). **Right:** MALDI-TOF MS. Calculated mass: 3229.714 Da $[M+H]^+$. Observed mass: 3228.641 Da $[M+H]^+$.

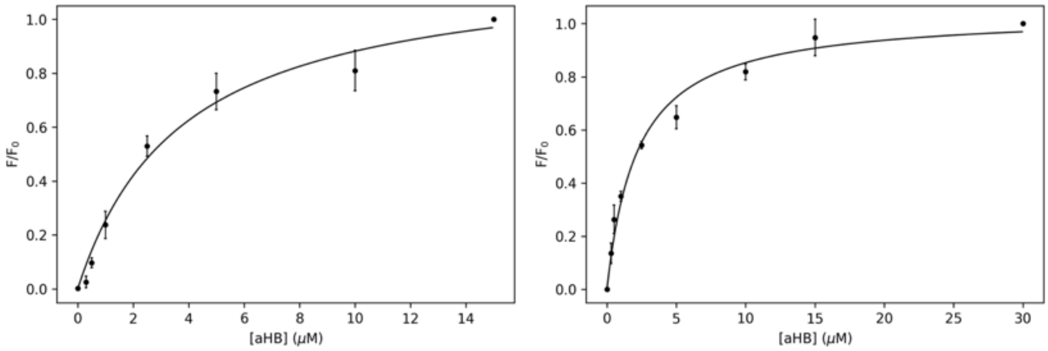

**Figure S9. Saturation binding curves with DPH. Left:** Saturation binding curve with DPH for CC-Type2-($S_gI_aI_d$)$_4$, returning a $K_D$ = 3.8 ± 0.8. **Right:** Saturation binding curve with DPH for CC-Type2-($A_gI_aI_d$)$_4$, returning a $K_D$ = 2.2 ± 0.3.

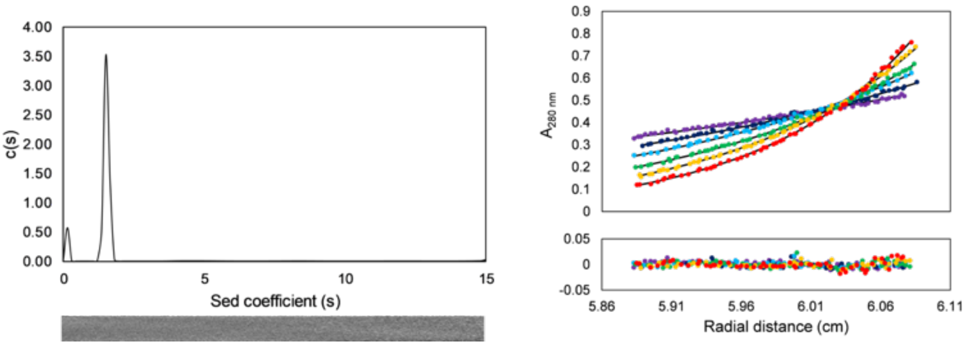

**Figure S10. Left:** Sedimentation velocity (SV) AUC data for CC-Type2-($T_gL_aI_d$)$_4$ at 50 krpm ($\bar{v}$ = 0.771 $cm^3 g^{-1}$). Residuals are shown as a bitmap below the fitted data. Continuous c(s) distribution returned a molecular mass of 16079 Da corresponding to 4.8 x monomer mass at 95% confidence level ($f/f_0$ = 1.19, s = 1.540 S, $s_{20,w}$ = 1.606 S). **Right:** Sedimentation equilibrium (SE) AUC data for CC-Type2-($L_aI_dG_e$)$_4$ between 15 and 30 krpm at 3 krpm intervals. Fitted single-ideal species model curves are overlaid in black and gave a molecular mass 16130 Da corresponding to 4.8 x monomer mass, 95% confidence limits 15980-16274 Da. Conditions: 150 μM and 70 μM peptide for SV and SE experiments respectively, PBS, pH 7.4, 20 °C.

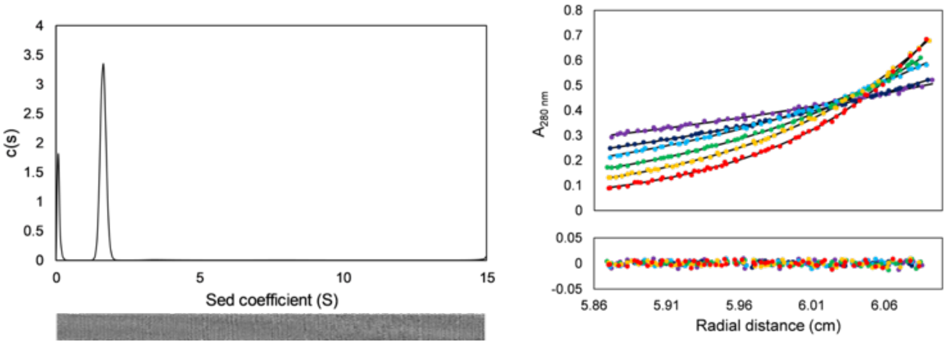

**Figure S11. Left:** Sedimentation velocity (SV) AUC data for CC-Type2-($T_gI_aI_d$)$_4$ at 50 krpm ($\bar{v}$ = 0.771 $cm^3 g^{-1}$). Residuals are shown as a bitmap below the fitted data. Continuous c(s) distribution returned a molecular mass of 18094 Da corresponding to 5.4 x monomer mass at 95% confidence level ($f/f_0$ = 1.21, s = 1.645 S, $s_{20,w}$ = 1.716 S). **Right:** Sedimentation equilibrium (SE) AUC data for CC-Type2-($L_aI_dG_e$)$_4$ between 15 and 30 krpm at 3 krpm intervals. Fitted single-ideal species model curves are overlaid in black and gave a molecular mass 17628 Da corresponding to 5.2 x monomer mass, 95% confidence limits 17475-17787 Da. Conditions: 150 μM and 70 μM peptide for SV and SE experiments respectively, PBS, pH 7.4, 20 °C.

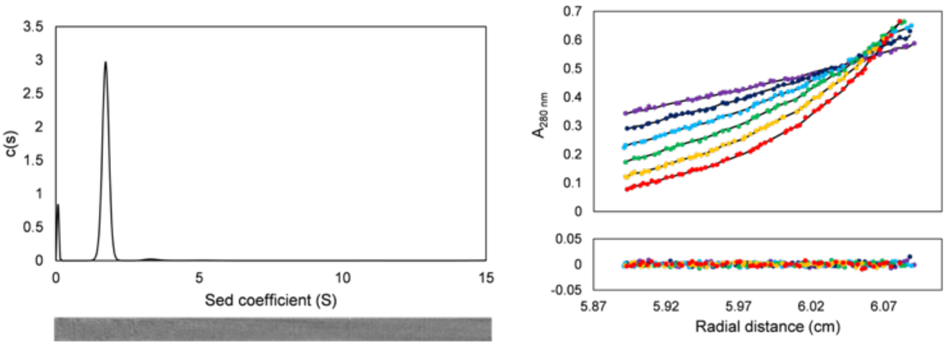

**Figure S12. Left:** Sedimentation velocity (SV) AUC data for CC-Type2-$(G_gL_aI_d)_4$ at 50 krpm ($\bar{v}$ = 0.771 cm$^3$ g$^{-1}$). Residuals are shown as a bitmap below the fitted data. Continuous c(s) distribution returned a molecular mass of 20890 Da corresponding to 6.5 x monomer mass at 95% confidence level ($f/f_0$ = 1.26, s = 1.739 S, $s_{20,w}$ = 1.814 S). **Right:** Sedimentation equilibrium (SE) AUC data for CC-Type2-$(L_aI_dG_e)_4$ between 15 and 30 krpm at 3 krpm intervals. Fitted single-ideal species model curves are overlaid in black and gave a molecular mass 19643 Da corresponding to 6.5 x monomer mass, 95% confidence limits 19488-19805 Da. Conditions: 150 µM and 70 µM peptide for SV and SE experiments respectively, PBS, pH 7.4, 20 °C.

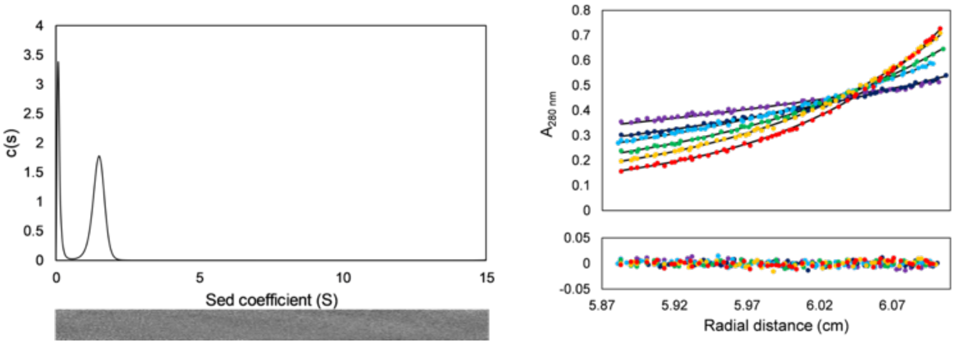

**Figure S13. Left:** Sedimentation velocity (SV) AUC data for CC-Type2-$(G_gI_aI_d)_4$ at 50 krpm ($\bar{v}$ = 0.771 cm$^3$ g$^{-1}$). Residuals are shown as a bitmap below the fitted data. Continuous c(s) distribution returned a molecular mass of 16050 Da corresponding to 5.0 x monomer mass at 95% confidence level ($f/f_0$ = 1.25, s = 1.473 S, $s_{20,w}$ = 1.536 S). **Right:** Sedimentation equilibrium (SE) AUC data for CC-Type2-$(L_aI_dG_e)_4$ between 15 and 30 krpm at 3 krpm intervals. Fitted single-ideal species model curves are overlaid in black and gave a molecular mass 16117 Da corresponding to 5.1 x monomer mass, 95% confidence limits 15931-16289 Da. Conditions: 150 µM and 70 µM peptide for SV and SE experiments respectively, PBS, pH 7.4, 20 °C.
